# Pyranges v1: a Python framework for ultrafast sequence interval operations

**DOI:** 10.64898/2025.12.11.693639

**Authors:** Endre Bakken Stovner, Max Ticó, Ester Muñoz del Campo, Joan Pallarès-Albanell, Konika Chawla, Pål Sætrom, Marco Mariotti

## Abstract

Sequence interval algebra is key to modern bioinformatics. Pyranges v1 offers a Python Pandas-based interface to a comprehensive palette of Rust-powered operations (e.g., overlap, count, slice intervals), enabling the intuitive development of efficient pipelines for diverse sequence data, including gene annotations, mapped reads, and protein domains. Pyranges is faster, consumes less memory, and offers more functionalities than alternative tools including BEDTools, emerging as an innovative one-stop shop for omics analysis.

## Text

Over the past two decades, advances in high-throughput “omics” technologies have revolutionized biology and biomedicine ^1–4^. Despite their disparate aims, most of these assays yield data that can be uniformly described as sequence intervals: start-end coordinates on DNA or protein sequences, often strand-specific and with additional quantitative and qualitative features. This abstraction unifies diverse biological data such as gene annotations, genetic variation, primers, sgRNA targets, protein domains, peptide hits, homology matches, and aligned reads from sequencing-based omics (RNAseq, Riboseq, ChIP-seq, ATAC-seq, Hi-C, methylation sequencing, and many others). Utilities for manipulation of sequence intervals, epitomized by BEDTools ^5,6^, constitute an essential foundation on which countless specialized pipelines are built. Omics workflows typically combine these elementary “genome arithmetic” operations (e.g. intersect, merge, complement, sort intervals) with conventional data tasks like filtering and aggregation.

Here we introduce Pyranges v1, a versatile Python library for ultrafast sequence interval analysis. V1 is a complete rewrite of our popular “v0” version ^7^ and it is tailored to the modern Python data-science stack. Each dataset is loaded in memory into a table-like PyRanges object wherein every row represents an interval. This data structure is now implemented as an explicit subclass of the DataFrame of Pandas ^8^, today’s most popular data analysis library. Users can thus directly tap into the rich and familiar Pandas API without incurring any conversion overhead, and seamlessly combine common data tasks with a broad palette of sequence interval operations offered by Pyranges (Figure 1). This includes a comprehensive suite of strand-aware overlap-related functions, such as intersect, merge, subtract, filter, cluster, overlap counting (Figure 1a) and many others (see Documentation online). Most functions accept a grouping identifier, thereby enabling intuitive transcript-level manipulation of multi-exon entities: users may elongate the 5’ and/or 3’ extremes of each mRNA; slice arbitrary portions (e.g. the first/last 100 nucleotides); or derive the outer limits of complete groups, e.g. genes (Figure 1b). Besides, Pyranges supports projections across hierarchical coordinate systems, allowing to map ranges bidirectionally between a local (e.g. transcript- or protein-based) and a global (genome-based) reference (Figure 1c). Fast reverse complement and translation operations on nucleotides are also offered via native methods, while fetching sequences from fasta files is provided via Pyfaidx ^9^. By combining these primitives, users can assemble sophisticated multi-omic workflows with just a few readable Python statements.

**Figure 1.**
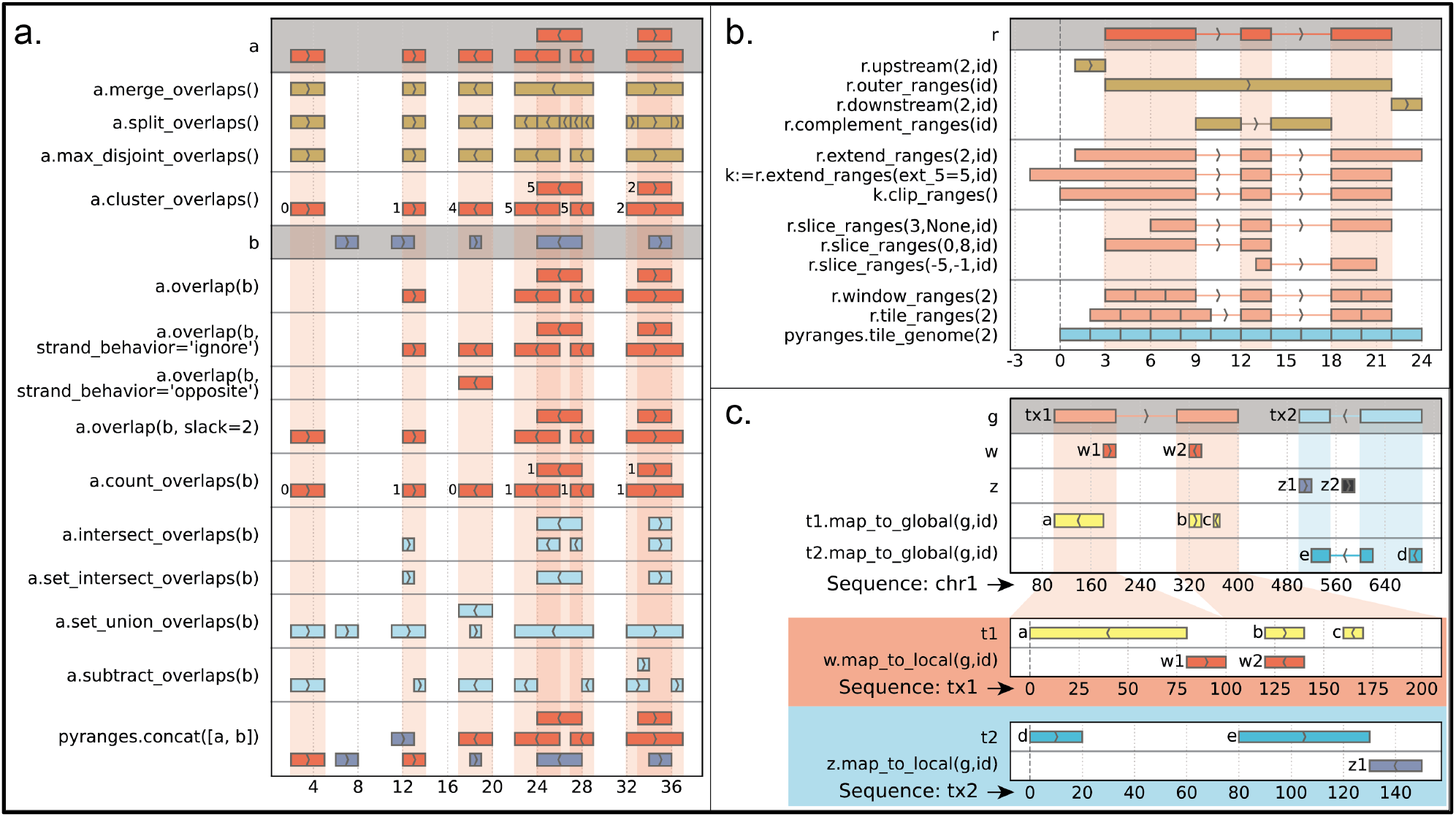
Cheatsheet showcasing some functions available in Pyranges. It includes (a) overlap-based operations, (b) single-object operations on coordinates, and (c) mapping across nested coordinate systems. Functions in (b) that include the “id” argument may act on groups of intervals, as shown here, or be applied to each interval independently. Derived data objects are shown with similar colors. Note that a single PyRanges object may host intervals mapped to multiple chromosomes, which is not showcased here. Refer to the online documentation for the full list of methods, including those not suitable for graphical representation (e.g. find nearest interval).

Beyond its user-friendly Python interface, Pyranges methods are implemented in Rust, granting high speed and memory efficiency. We compared its performance to three alternative tools for sequence interval operations: GenomicRanges (R library for in-memory data manipulation) ^10^, Bioframe (Python library, also in-memory) ^11^, and BEDTools (designed for text-stream processing) ^5,6^. We performed a comprehensive benchmark across common overlap-based operations: (i) find overlapping intervals; (ii) group them into clusters; calculate their union (“merge”) and (iv) difference (“subtract”); and (v) pair each interval with its nearest neighbor. We generated random intervals along two sequence sets (the human genome with 25 chromosomes, and the human proteome with 118,218 sequences) and we varied the maximum length of intervals (10, 100, 1000) and the total dataset size (10^4^, 10^5^, 10^6^, 10^7^). Our results show that Pyranges outperforms all alternative tools (Figure 2): on large datasets (≥10^6^ rows), Pyranges is on average 3.1, 5.4 and 15.7 times faster and 51%, 42% and 43% less memory-consuming than GenomicRanges, BEDTools, and Bioframe, respectively (full benchmark at Supplementary Figure S1, Supplementary Table T1). Our benchmark positions Pyranges as a versatile one-stop shop tool with top performance across all tasks, both with genome-scale coordinates and with the fragmented coordinate spaces of transcripts or proteins.

**Figure 2.**
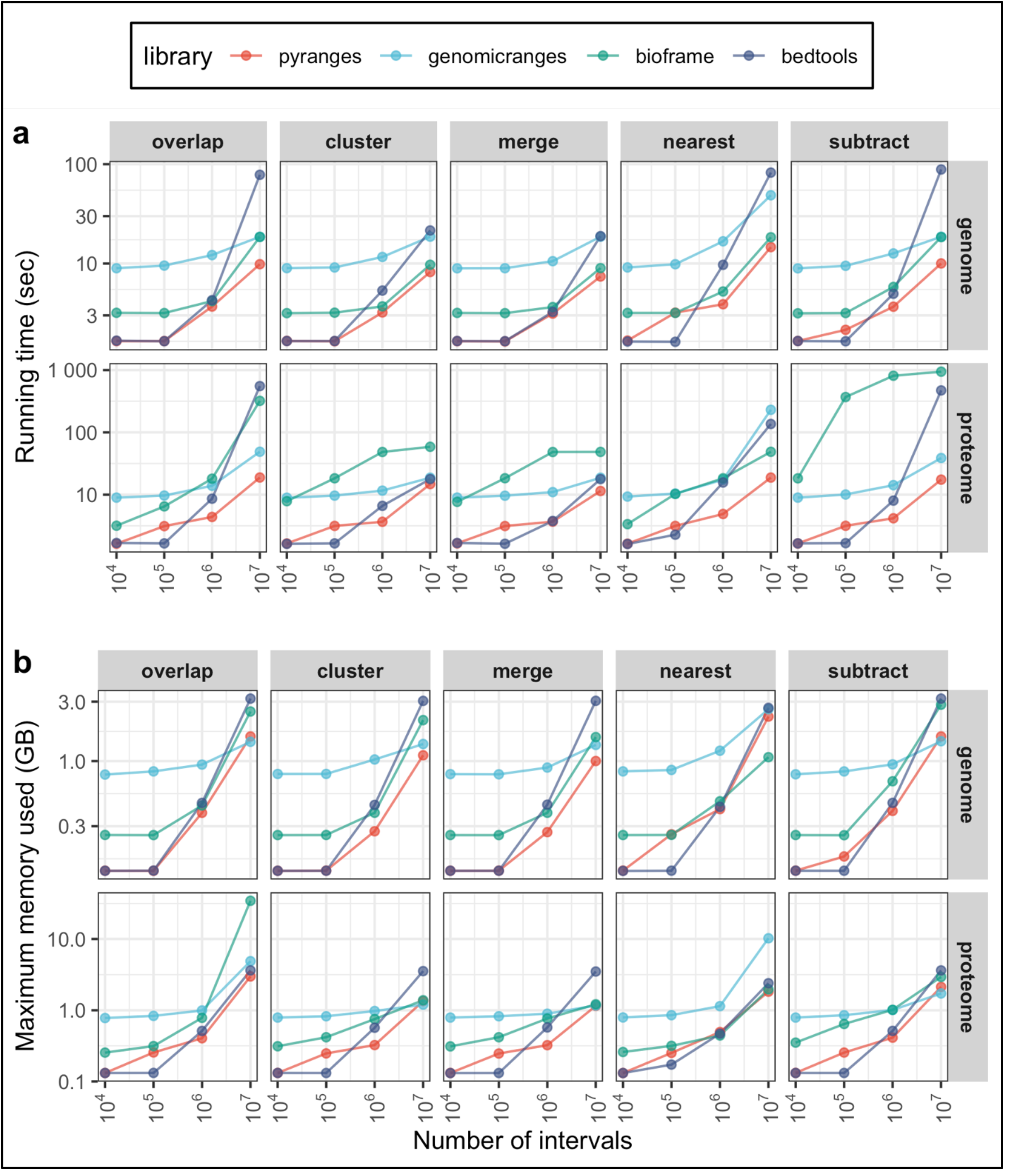
Benchmark of sequence interval tools’ speed (a) and memory usage (b). These results correspond to tests with the maximum length of intervals set to 100. Aggregate measures are presented in the main text. The complete benchmark results are available in Supplementary Figure S1 and Supplementary Table T1.

Notably, the workflows built with Pyranges are substantially shorter and easier to read than their counterparts with alternative tools, as evidenced in the benchmark notebooks themselves (Methods). Besides, Pyranges methods are arguably the most versatile, offering more options and behaviors. A comprehensive documentation (tutorial, how-to pages, and API reference) allows new users to rapidly learn the library’s capabilities and vocabulary. Moreover, Pyranges offers convenient methods to convert AI chatbots into specialized coding assistants, able to build complex workflows from simple user requests, as showcased in Supplementary Note 1.

Pyranges is designed to enable writing ultrafast genomic data pipelines with minimal, readable Python code. Besides, it is well-suited for data exploration within Jupyter notebooks and other interactive prompts, supported by convenient data access functions and intuitive representations, including an early prototype of the companion graphics library “Pyrangeyes”. These features establish Pyranges as an ideal tool for the modern, Python-oriented bioinformatician. Besides, to ensure accessibility beyond the Python ecosystem, the utility “Pyranger” offers access to the library’s core functionalities directly from the command-line, promoting its adoption by an even larger user base.

Pyranges welcomes contributions from a broad community, facilitated by a detailed guide for developers. Code quality and consistency are maintained through a robust continuous integration system on GitHub, which leverages the Python tool “tox” to run hundreds of tests across diverse environments for each new commit. Overall, this framework ensures that Pyranges will endure as a dependable toolkit in genomics for years to come.

## Supporting information

All supplementary files

## Online Methods

### Code and data availability

This paper refers to Pyranges v1.1.4, compatible with Python >=3.12. Source code: https://github.com/pyranges/pyranges_1.x (MIT).

Documentation: https://pyranges1.readthedocs.io.

Pyranges can be installed via pip (see https://pyranges1.readthedocs.io/installation/).

Pyranges builds upon the Ruranges library v0.0.13 written in Rust, available at https://github.com/pyranges/ruranges/.

### Benchmark

Benchmarking was implemented by adapting the Snakemake benchmark template. Benchmark scripts, as well as the code for plotting, are available at https://github.com/pyranges/benchmarks. The conditions tested are described in the main text. The data shown here was generated by running the Snakemake pipeline in single-core, single-job mode on a Linux server with an Intel Core i7 10700K / 3.8 GHz and 125GB of RAM. Comparatively similar results were obtained on a M2 MacBook Pro. While BEDTools provides a Python wrapper (pyBEDTools), we benchmarked against the faster, native command-line program. Versions of relevant programs/libraries used for benchmark: BEDTools v2.27.1, Bioframe v0.8.0, GenomicRanges v1.61.1, Pandas v2.3.0, Pyranges v1.1.2, Python v3.13.5, R v4.5.1.

### Graphics

Figures in this paper were generated with the Pyrangeyes prototype (v.0.1.1, available at https://github.com/pyranges/pyrangeyes), then adapted through PDF editing, and with R’s ggplot2.

#### Note for referees and editor

before publication, v1 will be merged into the default version (currently holding v0), and the links above will be updated.

## Acknowledgments

Authors thank all contributors to Pyranges code (https://github.com/pyranges/pyranges_1.x/graphs/contributors).

## Author contributions

E.B.S. wrote most of the software as well as the benchmarking framework. M.T., E.M.C., J.P.-A., and K.C. contributed to software development. P.S. provided feedback on results and improved the manuscript. M.M. contributed to software development, supervised the project, secured funding, prepared figures, and drafted the manuscript.

## Supplementary material

**Supplementary Table T1**. Complete benchmark results in Excel format. Three sheets with identical structure contain the benchmark results of three replicate runs, obtained as described in Methods.

**Supplementary Figure S1**. Complete benchmark results. Note that, in case of “fail” status (due to out-of-memory error), the time reported corresponds to the emergence of the error: the task was not completed.

**Supplementary Note 1**. AI-assisted prototyping of Pyranges workflows. This document showcases how users can turn a general use chatbot (in the example, ChatGPT) into a specialized coding assistant, able to produce Pyranges-based bioinformatics pipelines from requests written in plain English.

