## Supplementary material for "Pyranges v1: a Python framework for ultrafast sequence interval operations": All supplementary files: Supplementary Note 1.pdf

### Supplementary Note 1. AI-assisted prototyping of Pyranges workflows

We primed ChatGPT o3 with the prompt obtained by:

```
>>> import pyranges as pr
>>> pr.assistant.prompt(concise=True)
'Act as an expert bioinformatician programmer experienced in pyranges
(complete documentation attached for you to learn). Next, answer my
requests for code by first explaining the workflow, followed by
oneliner-style code snippets, as concise as possible but elegant,
preceded by the text of task as commented code. Ensure you use
pyranges v1 interface that you find here, rather than the v0, from
which you may have seen examples before; v1 renamed many methods.
Output code that is as concise as possible but elegant. Assume
pyranges is fully installed. No import statements. Use aptly named
variables, no need to declare them. '
```

Concomitantly uploading the file pyranges_docs.txt obtained with:

```
>>> pr.assistant.export_docs("pyranges_docs.txt")
```

We then posed five representative coding tasks. The code blocks below correspond to ChatGPT's first response, except when a NameError revealed an obsolete v0 method, in which case the traceback was provided and the model corrected the call on the next turn. No other edits or guidance were necessary, demonstrating that once primed with the v1 documentation, the assistant reliably delivered concise solutions that chain Pyranges primitives into complete workflows.

```
#1) file.gff has CDS entries. Make file.fasta with the first piece of each
encoded protein (10aa)
(pr.read_gff3('file.gff').query('Feature=="CDS"')
   .slice_ranges(0,30,group_by='ID')
   .get_sequence('genome.fa',group_by='ID')
   .pipe(pr.seqs.translate).str.slice(0,10)
   .pipe(lambda s: open('file.fasta','w').writelines(f'>{k}\n{v}\n' for k,v
in s.items())))
```

```
#2) file2.gtf has only exon and CDS entries. Derive and write 5UTR and 3UTR
coordinates to output.bed
((gt:=pr.read_gtf('file2.gtf')).query('Feature=="exon"')
.subtract_overlaps(gt.query('Feature=="CDS"'),
                   match_by='Parent',strand_behavior='same')
```

```
.to_bed('output.bed'))

#3) get all CDS regions from annotation.gff not covered by reads in file.bam
uncovered =
pr.read_gff3('annotation.gff').query('Feature=="CDS"').subtract_overlaps(pr.r
ead_bam('file.bam'))

#4) find the 10 genes from annotation.gff with most reads in the 1000bp
before their start
top10 = (pr.read_gff3('annotation.gff').query('Feature=="gene"')
           .upstream(1000)
           .count_overlaps(pr.read_bam('file.bam'))
           .sort_values('Count',ascending=False)
           .head(10))

#5) find the 20 genes with the shortest encoded proteins, write to table.tsv
(pr.read_gff3('annotation.gff').query('Feature=="CDS"')
   .groupby('Parent').apply(lambda d: ((d.End-d.Start).sum()//3))
   .nsmallest(20)
   .to_csv('table.tsv',sep='\t',header=False))
```
