## Supplementary figures and images for "Pyranges v1: a Python framework for ultrafast sequence interval operations"

### Supplementary Figure S1.pdf

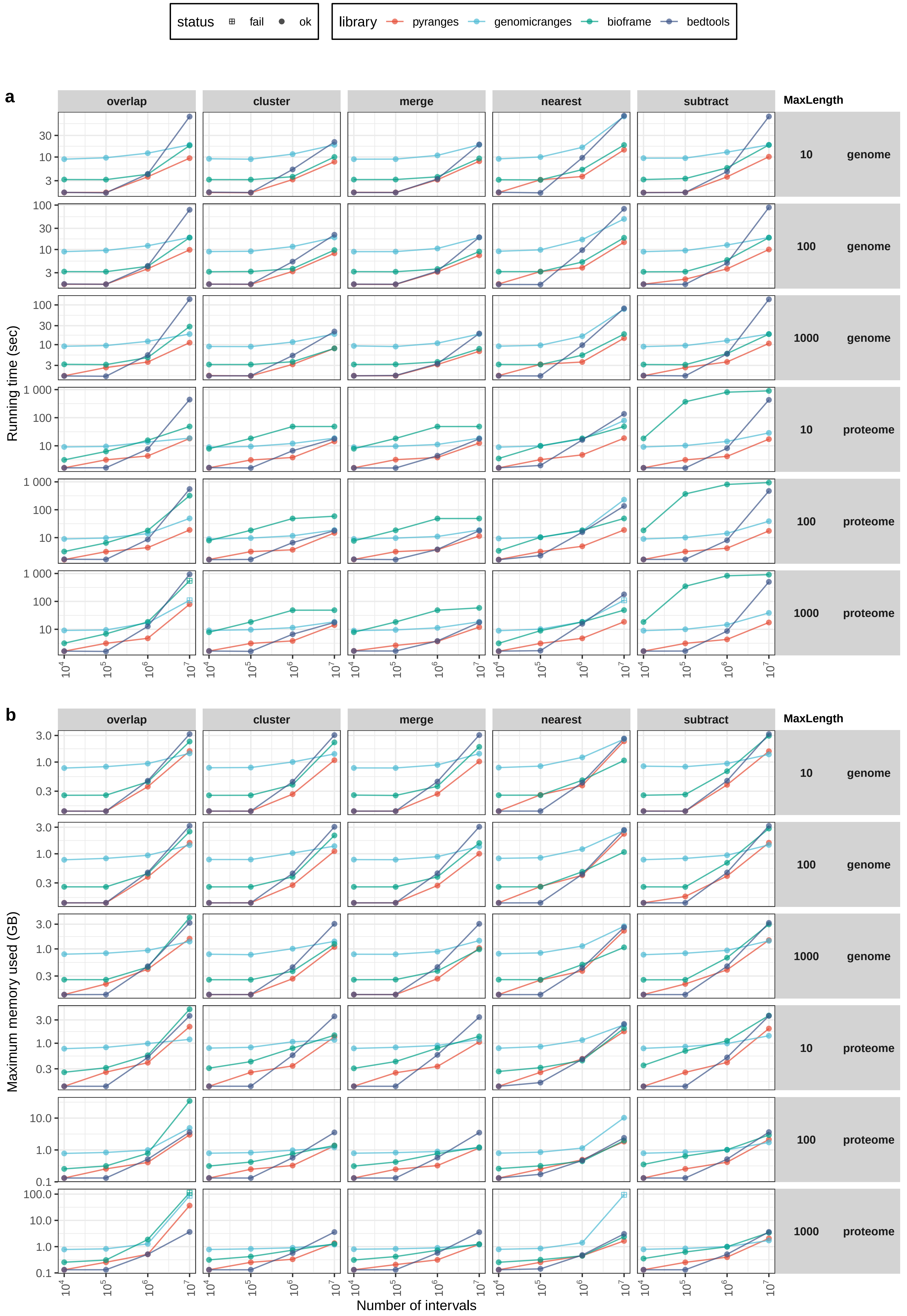
